# Oxidative stress drives mutagenesis through transcription coupled repair in bacteria

**DOI:** 10.1101/2022.06.28.497968

**Authors:** Juan Carvajal-Garcia, Ariana N. Samadpour, Angel J. Hernandez Viera, Houra Merrikh

**Affiliations:** Vanderbilt University School of Medicine, Department of Biochemistry, Nashville, TN 37232, USA; Department of Microbiology, University of Washington, Seattle, WA 98195, USA

## Abstract

In bacteria, mutations lead to the evolution of antibiotic resistance, which is one the main public health problems of the 21^st^ century. Therefore, determining which cellular processes most frequently contribute to mutagenesis, especially in cells that have not been exposed to exogenous DNA damage, is critical. Here, we show that endogenous oxidative stress is a key driver of mutagenesis and the subsequent development of antibiotic resistance. This is the case for all classes of antibiotics tested and across highly divergent species, including patient-derived strains. We show that the transcription-coupled repair pathway, which uses the nucleotide excision repair proteins (TC-NER), is responsible for endogenous oxidative stress-dependent mutagenesis and subsequent evolution. This strongly suggests that a majority of mutations arise through transcription-associated processes rather than the replication fork. In addition to determining that the NER proteins play a critical role in mutagenesis and evolution, we also identify the DNA polymerases responsible for this process. Our data strongly suggest that cooperation between three different mutagenic DNA polymerases, likely at the last step of TC-NER, is responsible for mutagenesis and evolution. Overall, our work identifies that a highly conserved pathway drives mutagenesis due to endogenous oxidative stress, which has broad implications for all diseases of evolution, including antibiotic resistance development.

## Introduction

Mutations provide the necessary genetic diversity that natural selection can then use to help organisms adapt to new environments. Even though mutations are mostly deleterious, and that lower mutation rates are generally accepted to be beneficial, all organisms have a baseline mutation rate that allows them to evolve (1). However, which mechanisms most commonly lead to mutations and drive evolution remain unknown.

An important source of mutations is damage to the DNA. However, the endogenous sources of DNA damage that cells are most commonly exposed to and lead to mutagenesis are unclear. This is partially due to the fact that most studies that investigate the mechanisms of mutagenesis in bacteria do so by exposing cells to high amounts of exogenous DNA damage, such as UV light. However, we reason that to better understand the mechanisms of evolution, we need an understanding of how cells that are not exposed to exogenous DNA damage mutate and evolve.

Oxidative stress is considered to be one of the main sources of endogenous DNA damage in bacteria. Oxidative stress is an obligatory consequence of aerobic respiration, and it results from an imbalance between highly reactive oxidative molecules (such as reactive oxygen species, ROS) and the cell’s ability to detoxify them (2). These oxidative molecules can react with biomolecules like proteins, lipids, and DNA, changing their chemical structure and damaging them. In the case of DNA, if this damage does not get properly repaired, oxidative stress can lead to mutations.

Accordingly, bacterial cells lacking catalase and superoxide dismutase, enzymes that de-toxify ROS, show growth defects as well as increased mutagenesis, even in the absence of exogenous DNA damage (3–5). Moreover, cells lacking glycosylases that excise oxidated DNA bases also show increased mutation rates (6, 7). We therefore hypothesize that endogenous oxidative stress plays a central role in bacterial evolution.

Here, utilizing antibiotics, we show that oxidative stress is the main source of mutations driving evolution. In addition, we determine that oxidative stress-dependent evolution is driven by nucleotide excision repair (NER), and in particular, transcription-coupled repair (TCR). We also show that a replicative polymerase, and two Y-family polymerases are responsible for the observed NER-dependent increase in mutagenesis. Critically, we show that all three polymerases function in the same pathway as the TC-NER proteins. Our results altogether show that a key source of mutations leading to evolution is oxidative stress induced TCR.

## Results

### Oxidative stress drives the evolution of antibiotic resistance

Oxidative stress has been proposed to be an important source of endogenous DNA damage in bacteria (2). For this reason, we considered whether decreasing the amount of oxidative stress bacterial cells are exposed to would have an effect on the kinetics of evolution. We utilized a previously described laboratory evolution assay to test this hypothesis (8). During this assay, we measured adaptation to the transcription inhibitor rifampicin in four different, highly divergent species: *Bacillus subtilis*, a multidrug resistant strain of *Staphylococcus aureus, Salmonella enterica* serovar Typhimurium and *Pseudomonas aeruginosa*. We have previously shown that the increase in the minimal inhibitory concentration (MIC) observed over time correlates with the appearance of mutations in known resistance genes (8, 9). We chose to test our hypothesis using rifampicin as it has been shown to not increase the amount of ROS in the bacterial cells (10), assuring that oxidative stress in the cells during our experiment has an endogenous origin.

We decreased the amount of oxidative stress in cells by adding the antioxidant thiourea, which has been used in the past to reduce oxidative stress in bacteria (10–12), at a concentration that does not affect the growth rate (Fig. S1A-D). In addition, to avoid antioxidant-independent effects of thiourea, we overexpressed *katA*, a gene coding for the ROS scavenging protein catalase, in *B. subtilis* (Fig. S1E) (13).

The staring MIC_50_ for the four species was 0.1 μg/ml (*B. subtilis*) 3.125 ng/ml (*S. aureus*), 8 μg/ml (*S. enterica*), and 4 μg/ml (*P. aeruginosa*) of rifampicin. (Fig. 1). After 35-40 generations in culture, the median concentration of antibiotic cells were able to survive increased to 3.2 μg/ml, 820 μg/ml, 1024 μg/ml, and 512 μg/ml of rifampicin respectively (Fig. 1). This value was significantly lower in cells that had been exposed to thiourea: 0.3 μg/ml (*B. subtilis*) 26.4 μg/ml (*S. aureus*), 256 μg/ml (*S. enterica*), and 20 μg/ml (*P. aeruginosa*) of rifampicin. Overexpression of *katA* in *B. subtilis* had a similar effect, as the median MIC on the last day of the experiment was 0.2 μg/ml of rifampicin (Fig. 1A).

**Figure 1:**
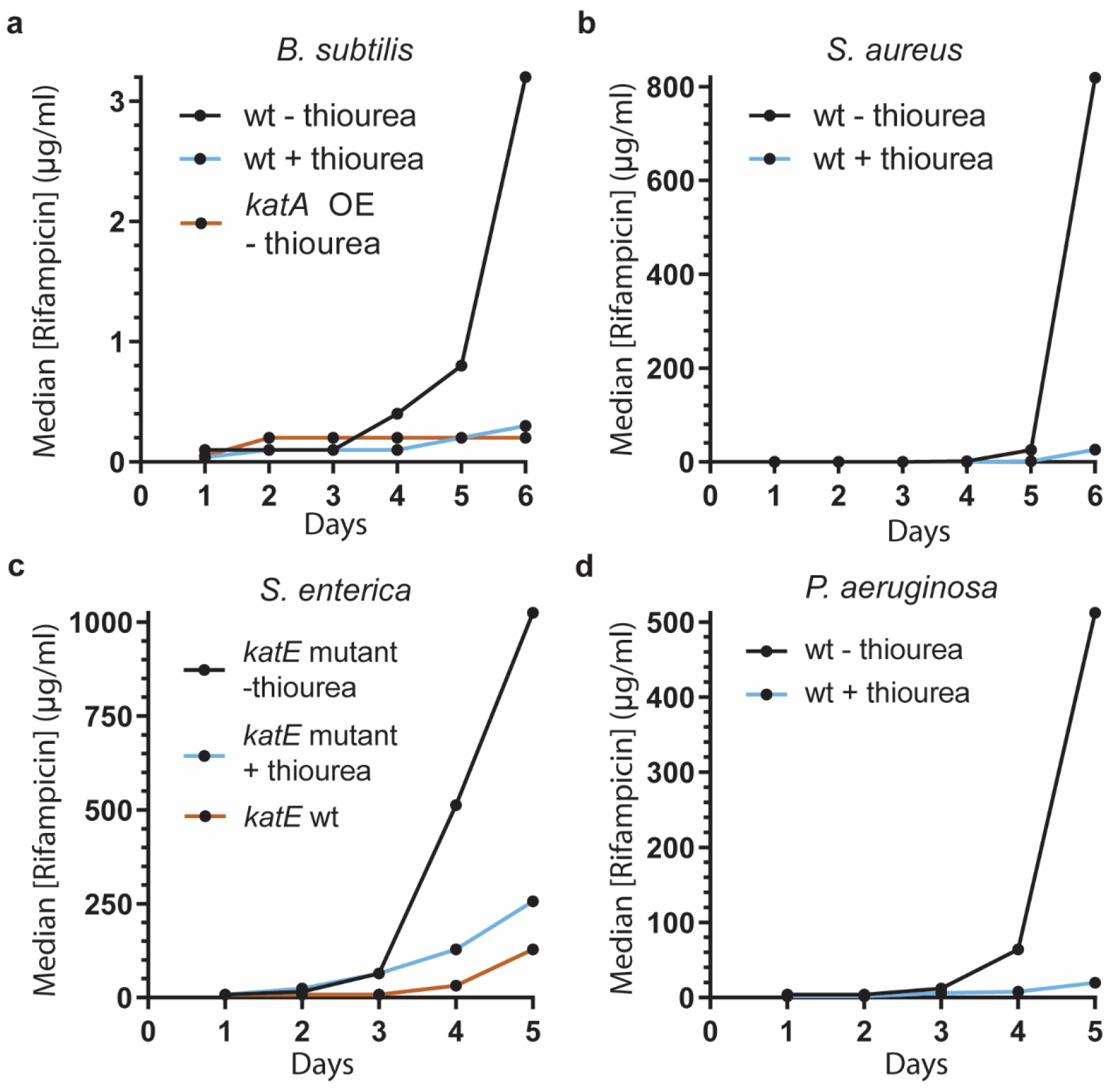
Oxidative stress drives the evolution of antibiotic resistance. Median concentration of rifampicin that allows for growth in the indicated strains at each sampled timepoint. 50 mM (a-c) or 10 mM (d) thiourea was included in the media where indicated. 1mM IPTG was added for *katA* overexpression. n=23 (*B. subtilis* – thiourea, rifampicin), 12 (*B. subtilis* + thiourea, rifampicin), 24 (*B. subtilis katA* overexpression, rifampicin), 12 (*S. aureus* – thiourea), 12 (*S. aureus* + thiourea), 35 (*S. enterica* serovar Typhimurium *katE* null – thiourea), 34 (*S. enterica* serovar Typhimurium *katE* null + thiourea), 24 (*S. enterica* serovar Typhimurium *katE* wt), 22 (*P. aeruginosa* – thiourea), 12 (*P. aeruginosa* + thiourea) biological replicates.

Interestingly, inactivating mutations in a catalase gene (*katE*) have been found in patient-derived *S. enterica* serovar Typhimurium strains (14), and this is the case with the strain that we used (ST19) (15). When we performed the same assay in a strain with a functional *katE* protein (SL1344) (15), we observed a reduction in the kinetics of evolution, similar to the one seen when adding thiourea (Fig. 1C). These observations are consistent with endogenous oxidative stress driving evolution generally in bacteria.

To test whether this phenomenon is universally conserved and not unique to rifampicin, we performed evolution assays using the translation inhibitor kanamycin and the folate synthesis inhibitor trimethoprim in *B. subtilis* as well as kanamycin and the cell wall synthesis inhibitor phosphomycin in *S. aureus*, and we obtained similar results (Fig. S1F-I).

### TCR drives oxidative stress-dependent evolution

We next decided to determine the mechanism by which endogenous oxidative stress drives evolution, using the genetically tractable species *B. subtilis*. Because oxidative DNA damage is commonly repaired by base excision repair (BER), we first tested whether BER mutants have decreased mutation rates, which would correlate with slower evolution of resistance. However, and consistent with previous reports (16, 17), we observed that strains lacking the DNA glycosylases MutY and MutM have higher mutation rates than wild-type cells (Fig. S2A).

We and others have previously shown that, in the absence of exogenous DNA damage, the bacterial TCR protein Mfd promotes mutagenesis across many different bacterial species (8, 18, 19). In addition, we previously showed that this pro-mutagenic effect depends on the interaction of Mfd with the RNA polymerase (RNAP) and the NER protein UvrA (8). Therefore, we decided to focus on nucleotide excision repair (NER). This DNA repair pathway has been shown to cause spontaneous mutagenesis in some bacteria (20–22), even if it has a protective effect against mutations when bacteria are exposed to DNA damaging agents (23, 24).

Bacterial NER has traditionally been described as consisting of two sub-pathways, global genome repair (GGR) and transcription coupled repair (TCR), differing in the damage recognition step (25). In GGR, UvrA scans the genome and binds DNA to trigger NER, and in TCR it is a stalled RNA polymerase who recruits the NER machinery to the site of DNA damage. This model has been put into question by recent studies claiming that, in bacteria, all NER is coupled to transcription (26, 27).

We performed an evolution assay in wild-type *B. subtilis* cells and in isogenic strains cells lacking the core component of the NER machinery UvrA, and we observed that UvrA promotes the evolution of antibiotic resistance (Fig 2A). In addition we measured mutation rates using the Luria-Delbruck fluctuation assay (28) in wild-type and mutants for the core NER proteins UvrA, UvrB and UvrC in the absence of exogenous DNA damage. We observed a 50-75% decrease in the mutation rates in NER deficient strains (Fig. 2B), indicating that NER also promotes spontaneous mutagenesis in *B. subtilis*.

**Figure 2:**
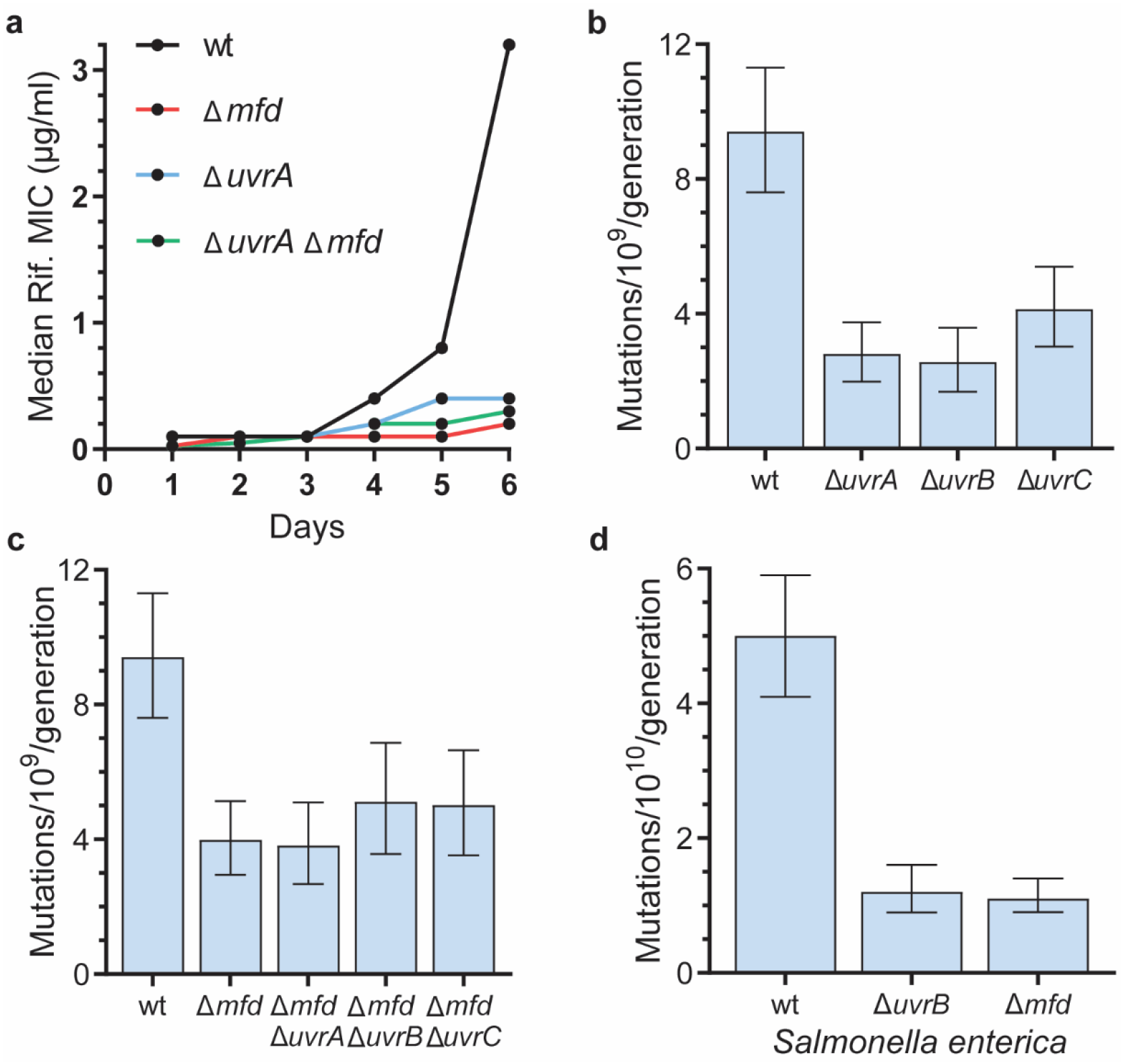
Transcription-coupled repair promotes mutagenesis. **a)** Median rifampicin concentration that allows for growth in the indicated strains at the indicated timepoints. n=23 (wt), 36 (*ΔuvrA*) 24 (*Δmfd*), 12 (*ΔuvrA Δmfd*) biological replicates. **b, c)** Mutation rates of *B. subtilis* strains measured using rifampicin. n=54 (wt), 48 (*ΔuvrA*), 37 (*ΔuvrB*), 48 (*ΔuvrC*), 59 (*Δmfd*), 40 (*Δmfd ΔuvrA*), 40 (*Δmfd ΔuvrB*), 50 (*Δmfd ΔuvrC*) biological replicates. **c)** Mutation rates of *S’. enterica* serovar Typhimurium strains measured using rifampicin. n=54 (wt), 40 (Δ*uvrB*), 48 (*Δmfd*). Error bars are 95% confidence intervals.

To test whether TCR was solely responsible for NER-dependent mutagenesis, we built double mutants that lacked Mfd and the Uvr proteins. If NER is mutagenic only due to TCR, then we expect that the double mutants lacking Mfd and NER proteins would have an epistatic relationship, and that the effect of the double mutants in mutagenesis and evolution would be similar to the single mutants. On the other hand, if NER-mediated mutagenesis is through both GGR and TCR, the combination of mutants lacking both Mfd and NER proteins would further reduce mutations. To discern between these possibilities, we performed evolution assays in *B. subtilis* cells lacking Mfd, as well as UvrA and Mfd both and observed a comparable decrease in the evolution of resistance in both single mutants and in the double mutant (Fig. 2A). In addition, we measured and compared the mutation rates of the single and double mutants side-by-side. We found that the mutation rates of strains lacking both Mfd and all three canonical NER factors have the same mutation rates as each single mutant alone (Fig. 2C). This strongly suggests that all (mutagenic) NER is coupled to transcription.

Additionally, we measured mutation rates in *S. enterica* serovar Typhimurium cells lacking either UvrB or Mfd, compare them to isogenic wild-type cells. We observed a similar result as in *B. subtilis* as, in the absence of either UvrB or Mfd, there was a similar decrease in mutation rates compared to a wild-type strain. These results suggest that the mutagenicity of NER being due to TCR is conserved amongst bacteria (Fig. 2D).

For both thiourea and *katA* overexpression, the decrease in the rate of evolution was similar to the one observed in strains lacking UvrA and/or Mfd (Fig. 1A, 2A). We therefore tested whether TCR is responsible for the mutagenic effect of endogenous oxidative damage, by performing evolution assays in strains deficient in TCR genes (Δ*uvrA* and Δ*mfd*) and adding thiourea. Although these strains have a serious deficiency in evolving resistance to antibiotics, towards the end of the evolution assays, a slight increase in their MIC can be observed (Fig. 2A). We took advantage of this and analyzed the rate at which evolution starts to take off at the last time points when oxidative stress is reduced. Consistent with our model, we observed that, in strains lacking UvrA or Mfd, thiourea did not have any effect on the rate of evolution (Fig. 3). This strongly suggests that TCR is driving mutagenesis and subsequent adaptive evolution dependent on endogenous oxidative stress.

**Figure 3:**
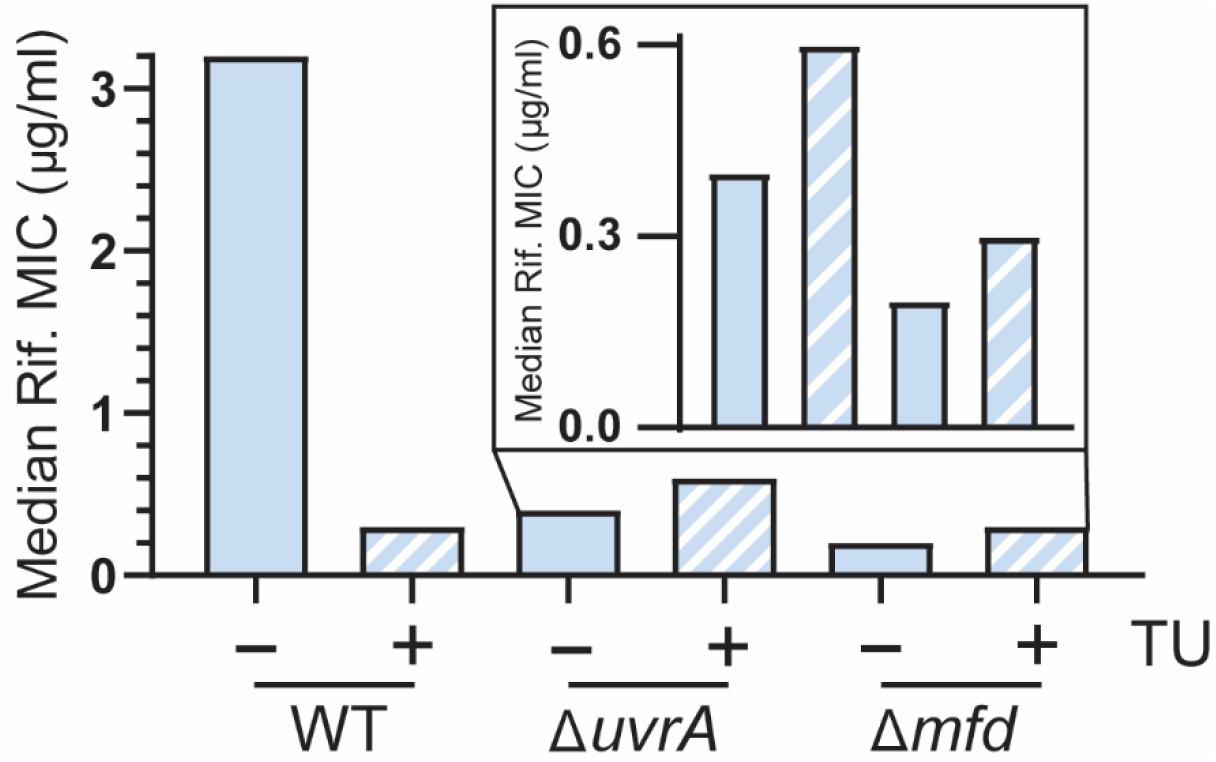
TCR promotes oxidative stress-dependent mutagenesis. Median rifampicin concentration that allows for growth in the indicated strains after six days of evolution. 50 mM thiourea was included in the media when indicated. n=23 (wt - thiourea), 12 (wt + thiourea), 34 (*ΔuvrA* - thiourea), 12 (*ΔuvrA* + thiourea), 24 (*Δmfd* - thiourea), 12 (*Δmfd* + thiourea) biological replicates.

### A replicative and two Y-family polymerases function in the same pathway as NER

We set out to determine the molecular mechanism behind NER-dependent mutagenesis. We reasoned that the gap filling step of NER is the most likely source of errors and that it may be completed by an error-prone mechanism. During NER, gap-filling synthesis is the last step of the pathway and, based on *in vitro* experiments, DNA polymerase I (PolA in *B. subtilis*) is thought to perform this step (29, 30). Thus, we measured mutation rates in cells lacking PolA. Although *in vitro* work had led to the conclusion that PolA is a high-fidelity polymerase (31), we found that this is not the case *in vivo*. We observed that PolA is mutagenic, as cells lacking PolA showed a decrease in mutation rates that were very similar to that seen in NER deficient strains (Fig 4A).

**Figure 4:**
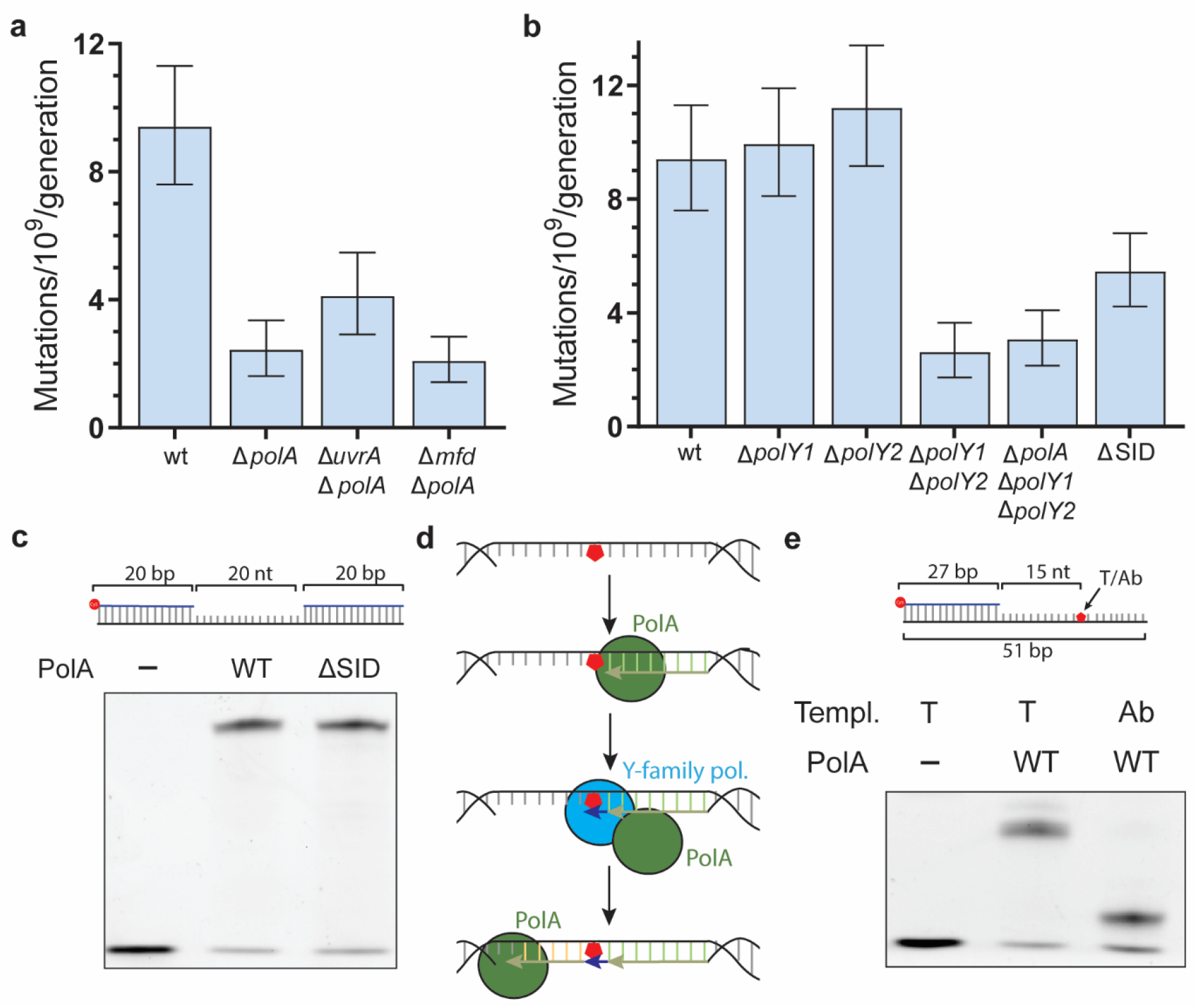
Three polymerases are required for TCR mutagenesis: **a, b)** Mutation rates of *B. subtilis* strains. n=54 (wt), 40 (*ΔpolA*), 36 (*ΔuvrA ΔpolA*), 44 (*Δmfd ΔpolA*), 57 (*ΔpolY1), 56 (ΔpolY2*), 35 (*ΔpolY1 ΔpolY2*), 43 (*ΔpolA ΔpolY1 ΔpolY2*) biological replicates. Error bars are 95% confidence intervals. **c)** Model for the molecular mechanism of NER-dependent mutagenesis. Due to DNA being single stranded in the transcription bubble and/or during NER, the non-transcribed strand is prone to damage that stalls PolA and leads to the recruitment of Y-family polymerases, further increasing the possibility of acquiring a mutation.

To determine whether NER is mutagenic due to PolA activity, we measured the mutation rates of *uvrA polA* and *mfd polA* double knockouts. When we compared the mutation rates of strains lacking either PolA, Mfd, or UvrA alone to the double mutants that lacked Mfd and PolA as well as UvrA and PolA, we did not see an additional decrease in mutation rates, indicating that Mfd- associated, mutagenic NER is in the same pathway as PolA (Fig. 4A). In addition, we used a biochemical assay where we purified *B. subtilis* PolA and used an *in vitro* primer-extension assay on a ssDNA gap template similar to the one that would be generated during NER to examine whether *B. subtilis* PolA can fill in this gap. We indeed observed that *B. subtills* PolA is able to efficiently fill in this gap (Fig. 4C, S2B).

However, *in vitro* studies with the *E. coli* ortholog of PolA (PolI) have determined that it is a high-fidelity polymerase, making it unlikely that by itself, it would introduce an error in such a small gap as the one generated during NER (31). Given that previous work has suggested that *B. subtilis* PolA interacts with two error-prone, Y-family polymerases, PolY1 and PolY2 (orthologs of the *E. coli* PolIV and PolV and the mammalian Pol kappa and Pol eta) (32), we reasoned that these Y-family polymerases could also be involved in the pro-mutagenic nature of NER. To test our model, we generated strains that lacked either PolY1, PolY2, or both polymerases. When we determined the mutation rates of strains that either lacked PolY1 or PolY2, we did not observe a decrease in mutation rates in either single mutant. Interestingly we did observe a decrease in mutation rates in strains lacking both PolY1 and PolY2, suggesting a redundant, pro-mutagenic role for these polymerases (Fig 4B).

If our hypothesis that these polymerases cooperate with PolA during the NER gap filling step is correct, then in strains that lack all three polymerases, we should not observe any additional decrease in mutation rates. Indeed, we observed that there was no additional decrease in mutation rates when cells lacked all three polymerases compared to cells either lacking only PolA, the Uvr proteins, or both PolY1 PolY2 (Fig. 4B, S2C). Therefore, we conclude that these polymerases are in the same pathway and cooperate to complete the last step during NER. The observed requirement for both an A-family replicative polymerase and a Y-family polymerase led us to the model outlined in Fig. 4D, in which PolA performs DNA synthesis during NER but will stall if a DNA lesion is present on the non-transcribed, NER template strand. This stalled PolA would then recruit a Y-family polymerase to overcome the lesion, increasing the chances of generating a mutation. The secondary lesion on the non-transcribed strand could occur when this region of the genome is single stranded during transcription or NER, which would render it more susceptible to damage (33).

To test this model, we created a 10 amino acid deletion in the endogenous *polA* gene that includes the predicted region of interaction between PolA and PolY1/2 (specific interaction domain, SID) (32), and observed that it leads to a decrease in mutation rates compared to wild-type cells (Fig. 4C). However, this decrease is smaller than a full *polA* deletion, which suggests that we are not destroying the interaction completely. Critically, we purified the PolA-ΔSID mutant and observed no difference with the wild-type protein in its ability to synthesize DNA using a ssDNA gap substrate (Fig. 4C, S2B), indicating that this decrease is not due to loss of PolA synthesis ability. In addition, using purified wild-type PolA, we measured DNA synthesis on an ssDNA template containing an abasic site, one the most common lesions observed in DNA (34), and a substrate for PolV in *E. coli* (35). We observed that this form of DNA damage is a strong block to synthesis by PolA (Fig. 5E), supporting the model that PolA alone cannot fill in a gap generated during NER if there is damage to the non-transcribed strand. This further supports our model for the involvement of both the high fidelity PolA and the low fidelity Y-family polymerases in the gap-filling step of NER.

## Discussion

Mutations generate the genetic diversity that evolution requires. Damage to the DNA is an important source of mutations, and since most organisms are not exposed to exogenous DNA damage, endogenous sources of damage likely plays a key role in evolution. In this work, we identified oxidative stress as the main source of endogenous DNA damage leading to mutagenesis and evolution in bacteria. Reactive oxygen species have been shown to lead to most spontaneous mutagenesis in *E. coli* (36) and are thought to be an important source of endogenous DNA damage (2). We tested the contribution of oxidative stress to mutagenesis by measuring the evolution of resistance to antibiotics in cells that have reduced oxidative stress, by either growing the in the presence of thiourea or overexpressing the catalase gene *katA*. We observed that the evolution of antibiotic resistance was slower in these conditions in diverse bacteria, including patient-derived strains (Fig. 1). Sublethal concentrations of certain antibiotics have been shown to lead to oxidative stress, and this phenomenon has been proposed to lead to antibiotic resistance (10, 11). However, we observe that oxidative stress leads to evolution of resistance to an antibiotic that does not seem to increase the production of reactive oxidative as well, such as rifampicin (10).

We then show that nucleotide excision repair (NER), which strongly suppresses mutagenesis in cells exposed to DNA damaging agents (23, 24), is actually promoting mutagenesis under endogenous conditions and is generally a pro-mutagenic mechanism. Bacteria lacking any one of the three core components of the NER mechanism, UvrABC, have lower mutation rates than wild-type cells, indicating that NER causes spontaneous mutations (Fig. 2). In addition, our data indicate that in the absence of exogenous damage, all NER functions in the same pathway as the transcription-coupled (TC) NER factor Mfd, suggesting that NER is universally transcription-dependent (Fig. 2). This is consistent with recent biochemical findings which suggest all bacterial NER is transcription-dependent (26). Interestingly, the little evolution we observe in NER deficient strains is *not* diminished in the presence of thiourea (Fig. 3), suggesting that oxidative stress-driven evolution is dependent on NER.

Our data shows that it is the cooperation of at least two DNA polymerases that causes NER-dependent mutations: the replicative polymerase commonly associated with NER, PolA, and one of two redundant Y-family polymerases, PolY1 and PolY2 (Fig. 4). We propose that DNA damage in the NER template strand (the non-coding strand, Fig. 4D) explains this requirement for both DNA polymerases to complete NER, which will naturally lead to an increased likelihood of mutations being introduced into the synthesized DNA gap. A similar model has been proposed to explain observations of mutagenesis in prokaryotes and eukaryotes, but in the presence of exogenous DNA damage (37, 38), including in a recent pre-print that uses a mouse liver cancer model in which cells are exposed to high levels of the DNA damaging agent diethylnitrosamine (39). This suggests that this mechanism of mutagenesis is universally conserved.

In our model, the DNA lesion that leads to mutagenesis is independent of the lesion that triggered NER. We propose that it may be caused during transcription and/or the execution of a damaged oligonucleotide by NER. The non-transcribed strand stays as ssDNA for an extended period of time during both processes, and it is well-known that ssDNA is more prone to oxidative damage than dsDNA (40, 41). Alternatively, the NER machinery has been shown to excise non-damaged DNA *in vitro* (42) and transcription stimulates this process *in vivo* (43). These gratuitous repair events, even if much less efficient than excision of damaged DNA, are predicted to be a common phenomenon, as the amount of non-damaged DNA outweighs the amount of damaged DNA by several orders of magnitude in cells that are not exposed to exogenous DNA damaging agents (44). Moreover, Mfd has been found to be bound to DNA throughout the genome in the absence of exogenous DNA damage (45, 46), and it plays a role in transcription that is independent of its role in TC-NER (47). This constitutive association with DNA and RNAP could lead to excision, fill in synthesis, therefore increasing the likelihood of mutations being introduced into undamaged DNA, without the need for a DNA lesion present in close proximity and on the opposite strand.

Last, we have described in a recent pre-print a small molecule inhibitor of Mfd, which delays the evolution of antibiotic resistance in many different pathogenic bacterial species (9). By identifying other factors in the Mfd-dependent, pro-mutagenic pathway, we have expanded the potential targets for an anti-evolution drug that can be used to minimize antibiotic resistance generation during the treatment of infections in the clinic (9).

## Materials and Methods

### Bacterial culture

*Bacillus subtilis*, *Salmonella enterica* serovar Typhimurium, and *Staphylococcus aureus* were cultured in lysogeny broth (LB), and *Pseudomonas aeruginosa* in LB with 0.1% tween 20 (when liquid media). Bacterial plates were grown overnight at 37 °C unless otherwise indicated with the following antibiotics when appropriate: 500 μg/ml erythromycin and 12.5 mg/ml lincomycin (MLS), 5 μg/ml kanamycin (*B. subtilis*) or 50 μg/ml (*E. coli* and *S. enterica*) kanamycin, 25 μg/ml chloramphenicol and 100 μg/ml carbenicillin. When grown in liquid media, cultures were started from single colonies and were grown with aeration (260 rpm). A list of all strains used in this study can be found in Supplementary Information Table 1.

### Strain construction

The parental strain for all *B. subtilis* strains used in this study is HM1 (same as AG174, originally named JH642) (48, 49). Gene deletions that are marked with MLS or kanamycin resistance were obtained from (50). Genomic DNA from these strains was purified with the GeneJET Genomic DNA Purification Kit (Thermo) following the manufacturer’s instructions and transformed into the HM1 background as in previously described (51). When necessary to make strains that carry multiple mutations, these antibiotic resistant cassettes were excised by transforming the strains with a plasmid expressing the Cre recombinase (pDR244, BGSCID: ECE274) purified from RecA+ *Escherichia coli* (NEB) cells with the GeneJET Plasmid Miniprep Kit (Thermo), generating markerless strains (50). Recombinants containing markerless deletions were checked by PCR (Supplementary Information Table 2).

For *katA* overexpression, the coding sequence of the *katA* gene was amplified using Q5 polymerase (NEB) (Supplementary Information Table 2) and cloned between the *HindIII* and the *NheI* sites in pCAL838 (52) to form pHM724. pHM274 was linearized with *KpnI* and transformed into competent HM1 cells. Cells were plated on MLS containing plates and after overnight incubation at 37 °C, MLS resistant colonies were tested for growth in media lacking threonine. Colonies that lack growth in media without threonine and were MLS resistant were selected as double crossover integrants.

For deleting the specific interaction domain (SID) (32) of the endogenous *polA* gene in *B. subtilis*, 1 kb of homology on each side of the SID was amplified and cloned using NEBuilder® HiFi DNA Assembly Master Mix into the BamHI site of pMiniMAD2 (53), generating pHM736. This plasmid was transformed into RecA+ *E. coli* cells, purified, and transformed into HM1 *B. subtilis* cells, which were plated into MLS plates. After 48 hours at room temperature, a single colony was streaked in a fresh MLS plate and grown at 42 °C for 8 hours to force plasmid integration. A single colony was inoculated into liquid media and grown at 24 °C for 8 hours, then diluted 1:30 and grown at 24 °C for 16 hours, this was repeated for three days. Cells where then streaked on plates without antibiotics and grown at 37 °C overnight. Single colonies where then grown in plates with and without MLS at 42 °C for 8 hours. For colonies that are MLS sensitive, DNA was extracted and a PCR surrounding the SID was performed and ran on native PAGE gel. A colony with an apparent deletion was sequenced to confirm the expected 30 bp deletion.

The *S. enterica* Typhimurium strains ST19 and SL1344 (15, 54) were a gift from Sam Miller (University of Washington) and Mariana Byndloss (Vanderbilt University) respectively, the *Pseudomonas aeruginosa* strain is CF127 (55) and was a gift from Matt Parsek (University of Washington) and the multidrug-resistant *Staphylococcus aureus* strain is a cystic fibrosis patient derived strain obtained from the Vanderbilt University Medical Center.

For *S. enterica*, knock outs were made from the SL1344 strain by recombineering as previously described (56) using the pSIM27 plasmid, a gift from the Court lab (https://redrecombineering.ncifcrf.gov/strains--plasmids.html). In short, for knocking out *mfd*, the chloramphenicol resistance gene was amplified from the pKD3 plasmid (a gift from the Wanner lab (57)) while adding 40 nucleotides of homology upstream of the start site and downstream of the stop codon using Q5 polymerase (Supplementary Information Table 2). The PCR amplicon was cleaned and electroporated into competent, wild-type cells harboring the pSIM27 plasmid. Chloramphenicol resistant colonies were selected and checked by PCR (Supplementary Information Table 2). For knocking out *uvrB*, the kanamycin resistance gene was amplified from an *E. coli* strain with this gene on its chromosome (Supplementary Information Table 2) and recombineering was performed as described above.

### RNA extraction and gene expression levels determination

To measure the expression of the *katA* gene, single colonies were grown for 3 hours in 10 ml of LB to reach exponential phase. Cultures were then diluted an OD of 0.05 in 10 ml of LB including 1 mM IPTG and cultured for an hour (3 generations). RNA was extracted with the GeneJET RNA Purification Kit (Thermo) following the manufacturer’s instructions. 500 ng of RNA was treated with DNase I, RNase-free (Thermo), and cDNA was synthesized with the iScript™ cDNA Synthesis Kit (Biorad) using random primers in 20 μl reactions. Gene expression was determined by qPCR using the SsoAdvanced™ Universal SYBR® Green Supermix (Biorad) in 12 ul reactions, 5 μl of cDNA was used for the detection of *katA* and 2 μl for the detection of the 16S rRNA cDNA (used as housekeeping gene). Relative gene expression was calculated by the ΔΔCt method. For statistical comparison, ΔCt values were used.

### Evolution assays

Evolution assays were performed as previously described (8). A single colony of the indicated species and genotype was grown until and OD of 1-2 was reached. Culture was then diluted to and OD of 0.01 in culture media and grown in 7 different concentrations of the indicated antibiotic, ranging from no antibiotic to 16X the minimal inhibitory concentration (MIC), as well as thiourea or IPTG when indicated. Cells were grown for 24 hours at 37 °C with aeration, after which the OD was measured. The culture with the highest antibiotic concentration that showed an OD larger than 0.5X the OD of the culture without antibiotic (or, in the case of *P. aeruginosa*, an OD>0.3) was diluted 100X to an OD of approximately 0.01 and again grown in 7 different antibiotic concentrations. This process was repeated 6 times unless the fastest evolving strains reach an MIC higher than the solubility of the antibiotic in media.

### Determination of the mutation rates by fluctuation assays

Mutations rates were calculated as previously described (8). A single colony was inoculated into 2 ml of LB and grown for 2 hours (*B. subtilis*) or 2.5 hour (*S. enterica*) to reach exponential growth (0.1 <OD<0.6). This culture was diluted to an OD of 0.0005 and between 3 and 10 parallel cultures with 2 ml of LB were grown for 4.5 hours. Then, 1.5 ml of cells were pelleted and plated on 50 ug/ml rifampicin containing plates. The remaining cells were serially diluted in 1X Spizizen media and plated on antibiotic free media to quantify total viable cells. Colonies were counted after 24 hours at 37 °C (rifampicin plates) or 16 hours at 30 °C (no antibiotic plates). Mutation rates were calculated by using the Fluctuation AnaLysis CalculatOR (58), utilizing the Ma-Sandri-Sarkar maximum likelihood method.

### Growth curves

Growth curves were determined by growing a single colony of the indicated species until and OD of 1-2 was reached. The culture was diluted to an OD of 0.01 in culture media and growth in an Epoch microplate spectrophotometer (BioTek) at 37 °C for 16 hours. OD600 was measured every 10 mins.

### PolA purification

The coding sequence of PolA without the start codon was amplified by PCR using Q5 polymerase (NEB) and cloned BamHI-XhoI into pET28a (Thermo) to generate an N-terminal 6X his tagged protein coding sequence. The plasmid was transformed into BL21(DE) cells (NEB), and a single colony was inoculated into 70 ml of LB and grown overnight in LB containing kanamycin. 10 ml of culture were then inoculated in 1 L of LB+kanamycin and grown until an OD600 of 0.6, when 1 mM IPTG was added to the media. Cells were grown for 4 hours and centrifuged for 15 mins at 4000G. Pellets were resuspended in 30 ml of CelLytic B cell lysis reagent (Sigma) with 3 μl of Benzonase (Sigma) and 10 mM imidazole and shaken at RT for 10 mins. Lysate was centrifuged at 20000G at 4 °C and the supernatant was mixed with an equal volume of equilibration buffer (20 mM sodium phosphate pH 7.4, 300 mM sodium chloride, 10 mM imidazole), and run twice through 15 ml of equilibrated HisPur™ Ni-NTA Resin (Thermo) at 4 °C. Resin was washed with 150 ml of wash buffer (20 mM sodium phosphate pH 7.4, 300 mM sodium chloride, 40 mM imidazole) and eluted with 15 ml of elution buffer (20 mM sodium phosphate pH 7.4, 300 mM sodium chloride, 150 mM imidazole). Protein was dialyzed with a 30 ml Slide-A-Lyzer Dialysis Cassette G2 20000 MWCO (Thermo) against 10 mM tris pH 8, 50 mM NaCl, 5% glycerol, 0.1 mM DTT, 0.1 mM EDTA overnight at 4 °C Protein prep was then concentrated with Amicon Ultra-15 Centrifugal Filter Units 3000K (Millipore) to a final concentration of 1.6 mg/ml measured by Bradford assay (Thermo).

For the preparation of PolA-ΔSID, pET28a-PolA was used as template for PCR with 5’ phosphate containing primers, separated by the SIM sequence (amino acids 469-478) going outwards using Q5 polymerase (NEB). PCR product was purified, digested with DpnI (NEB) to eliminate the template plasmid, purified, and ligated using T4 DNA ligase for 1 hour at room temperature. Purification was performed as described above for the wild-type protein to a concentration of 2.1 mg/ml. Both protein preps were run on a 10% SDS-PAGE and stained by Imperial protein stain (Thermo) to confirm purity of purified enzyme.

### PolA synthesis assay

PolA synthesis was tested on 40 mM Tris pH 8, 10 mM MgCl2, 60 mM KCl, 2.5% glycerol buffer containing 1 mM dNTPs, 1.5 nM of the indicated DNA substrate labeled with Cy5, and 100 nM PolA. 10 ul reactions were incubated at 37 °C for 10 mins and stopped with 10 ul of 95% formamide 10 mM EDTA. DNA was denatured at 85 °C for 15 mins and run in a 12% urea denaturing gel at 150V for 30 mins. Gel was scanned in a ChemiDoc imaging system (BioRad).

The substrates for PolA synthesis experiments were done by annealing three (gap substrate) or two (primer extension substrate) HPLC purified oligos (Sigma) in a thermocycler. The template for the abasic site substrate contained a deoxyuracil in the 9th position. The abasic site was generated by treating the annealed oligo with hSMUG1 (NEB) for 30 mins at 37 °C followed by heat inactivation of the enzyme at 65 °C for 20 mins.

## Acknowledgements

We would like to thank current and former Merrikh Lab members for helpful discussion, especially Anna Johnson.

## Supplementary Information

**Supplementary Information Table 1:**
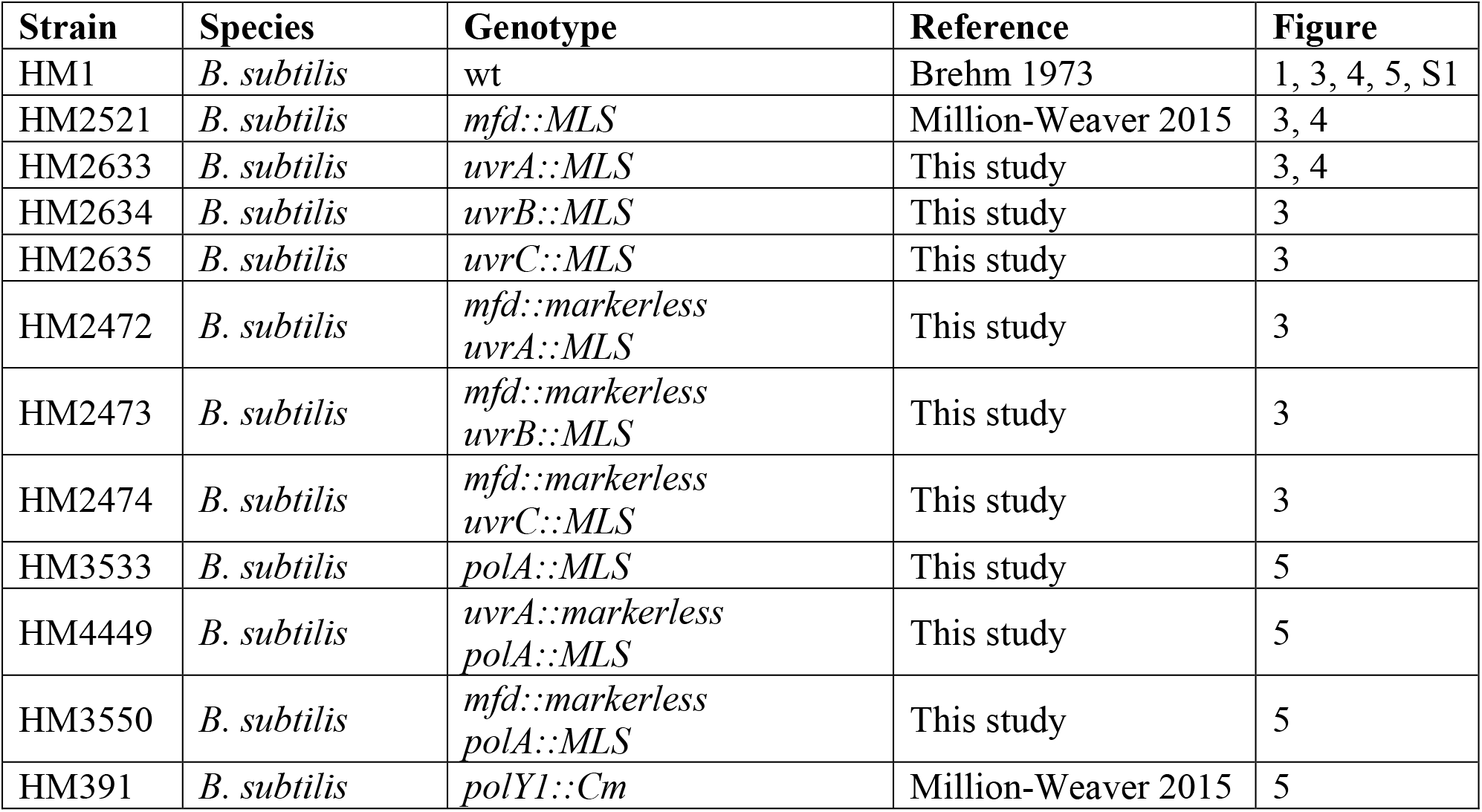
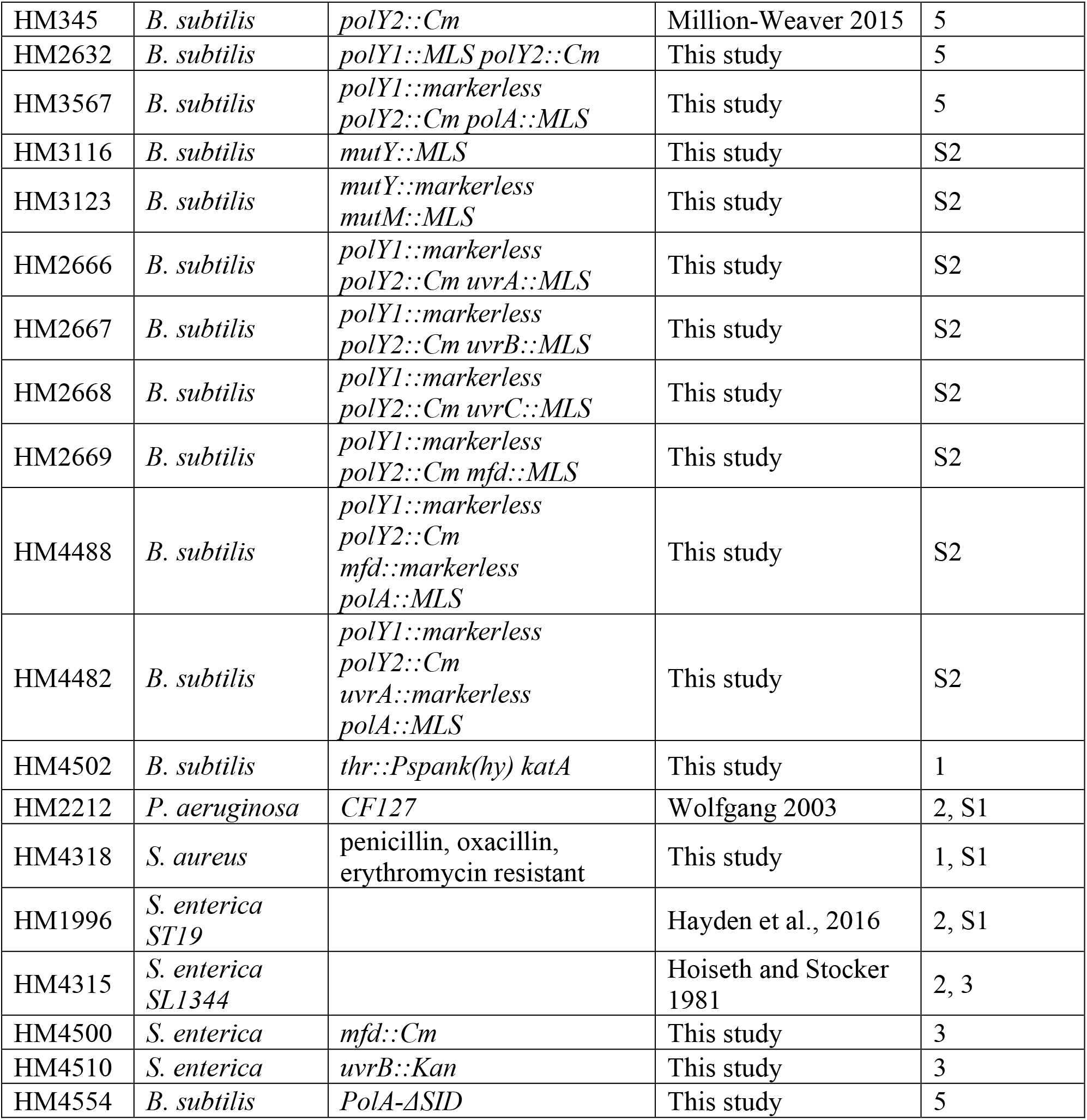
Strains used.

**Supplementary Information Table 2:**
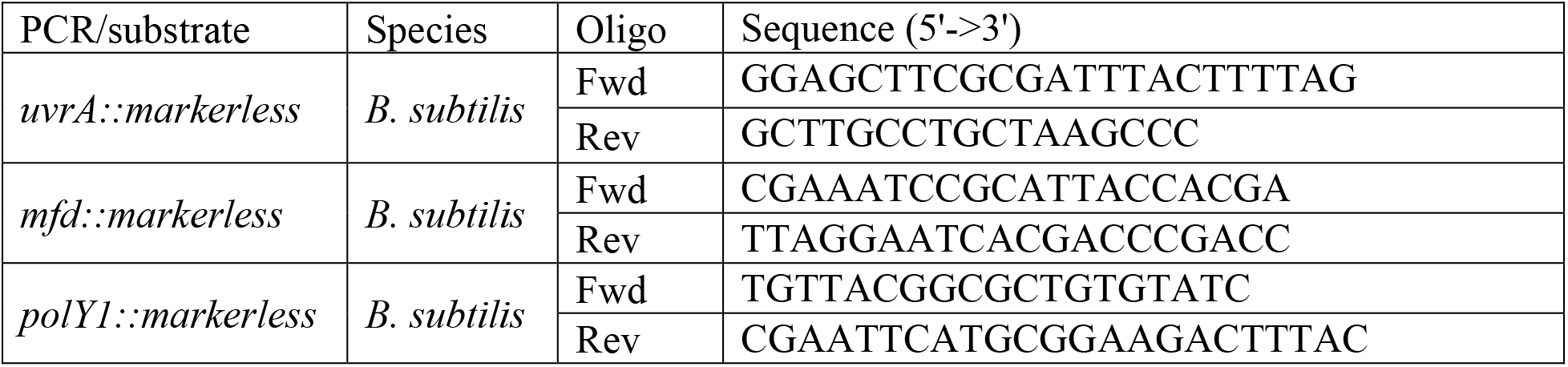
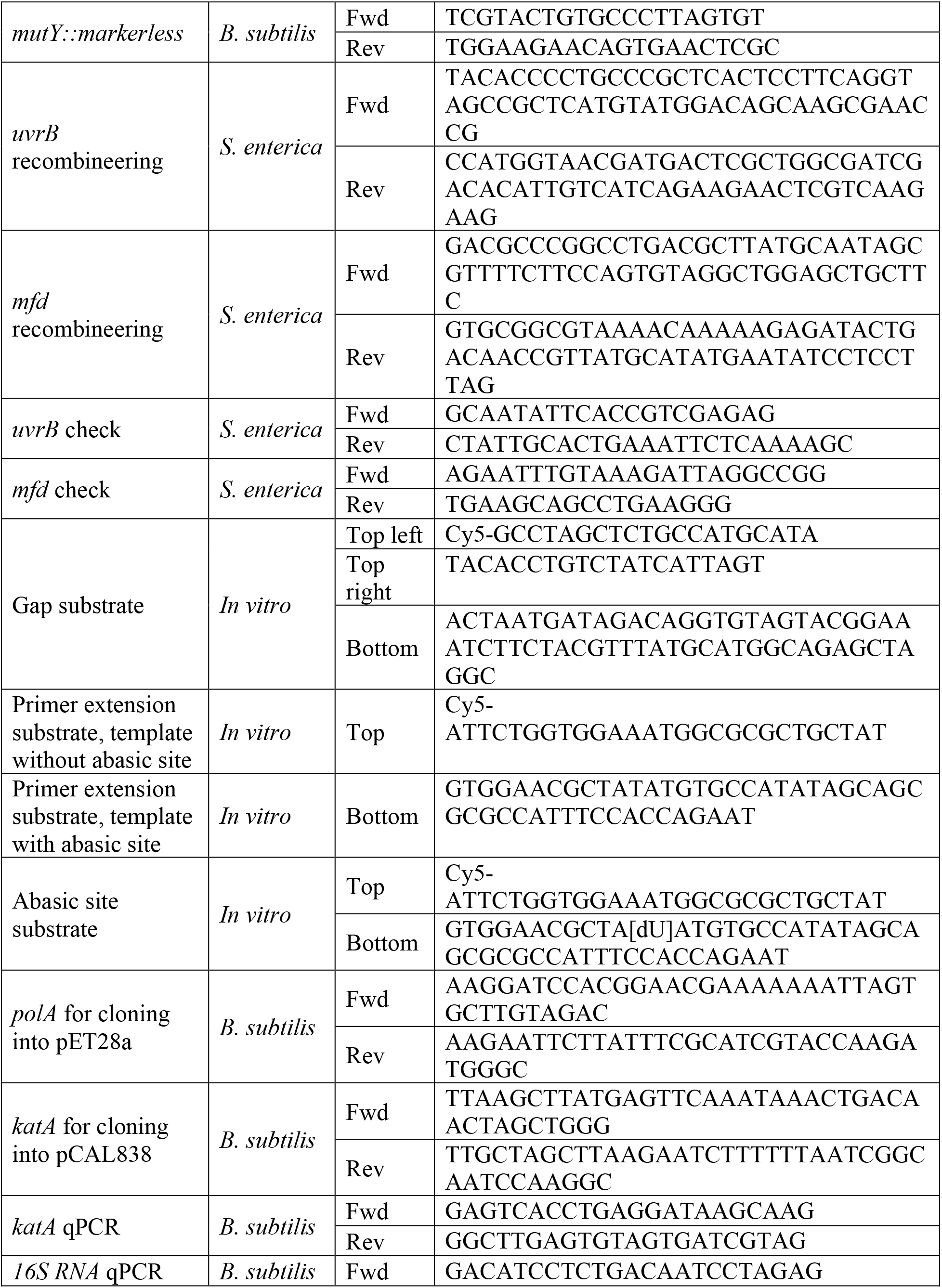
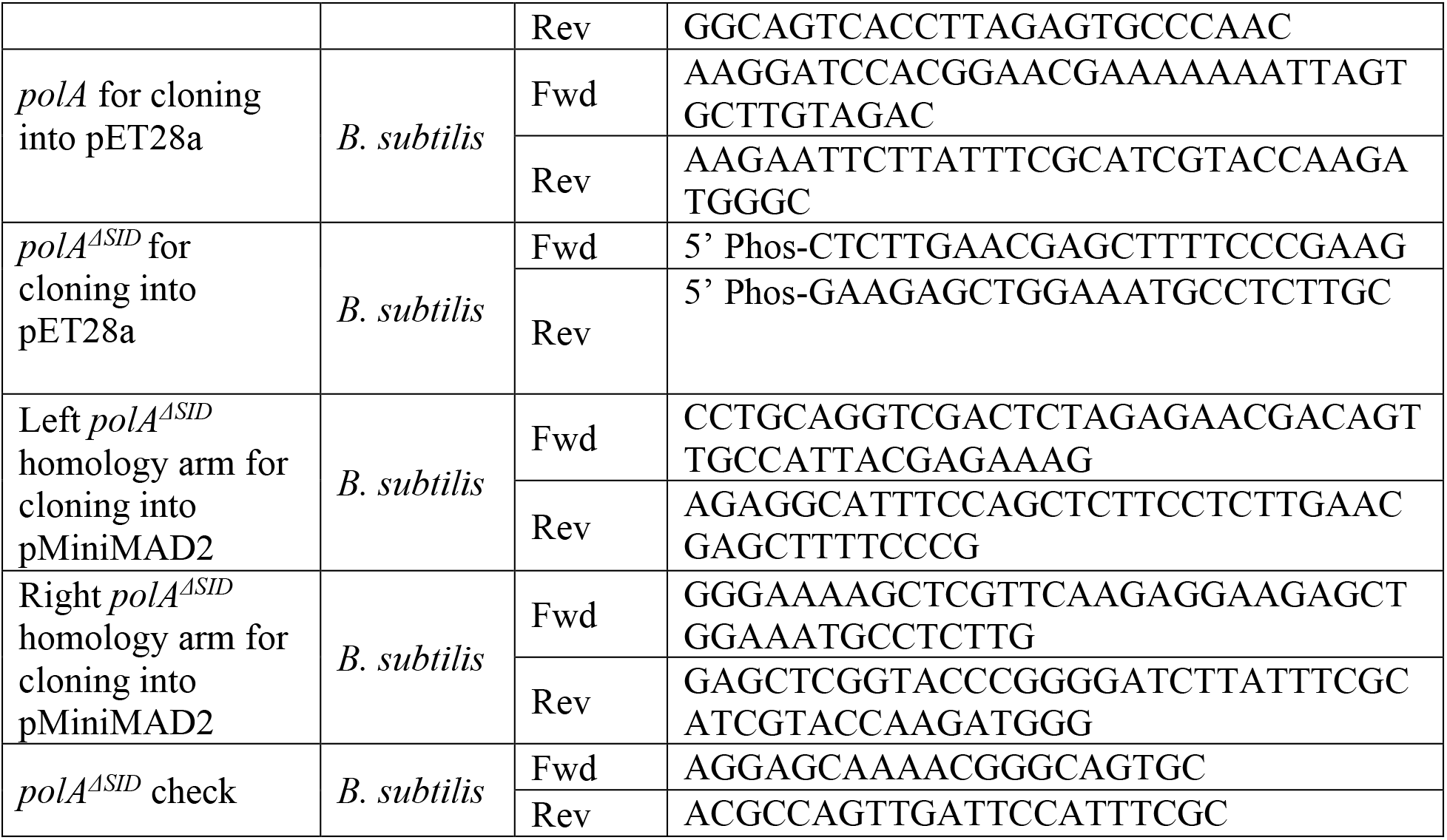
Oligonucleotides used.

**Supplementary Information Figure 1:**
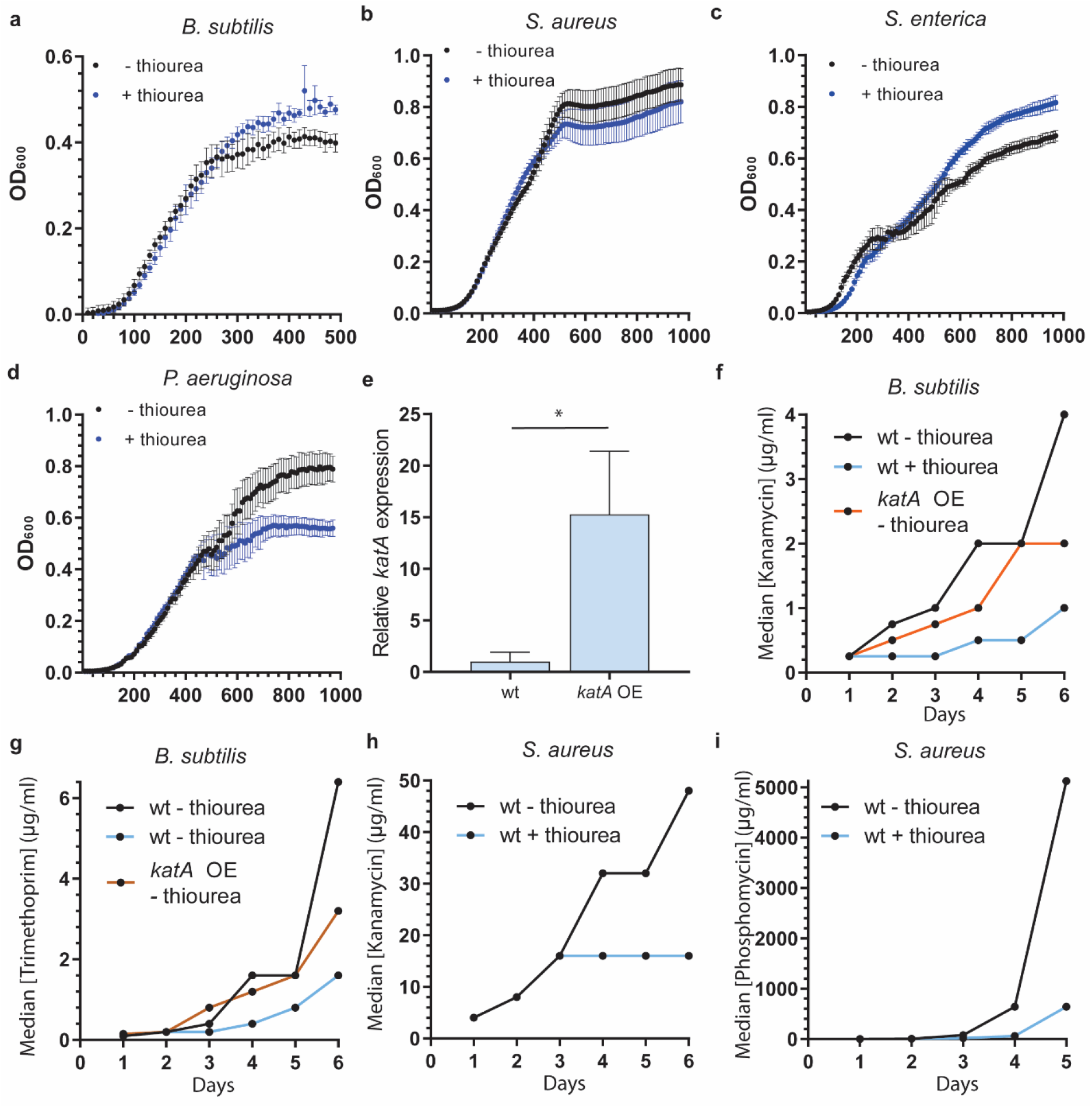
a-c) OD600 measured every 10 mins for the indicated time in a cultures of *B. subtilis* (a), *S. aureus* (b), and *S. enterica* serovar Typhimurium ST19 (c), with and without 50 mM thiourea in the media, n=12 biological replicates. d) OD600 measured every 10 mins for the indicated time in a culture of *P. aeruginosa* with and without 10 mM thiourea in the media, n=12 biological replicates. e) Normalized *katA* cDNA detected by qPCR in wild-type and *katA* overexpressing cells in the presence of 1 mM IPTG, n=5 biological replicates. f-i) Median concentration of rifampicin that allows for growth in the indicated strains at each sampled timepoint. 50 mM thiourea was included in the media where indicated. 1mM IPTG was added for *katA* overexpression. n=24 (wt – thiourea, kanamycin), 11 (wt + thiourea, kanamycin), 12 (*katA* overexpression, kanamycin), 12 (wt – thiourea, trimethoprim), 12 (wt + thiourea, trimethoprim), 12 (*katA* overexpression, trimethoprim), and 12 for all *S. aureus* experiments. Error bars indicate standard deviation. Statistical significance was assessed with two-tailed t-test, *p<0.05)

**Supplementary Information Figure 2:**
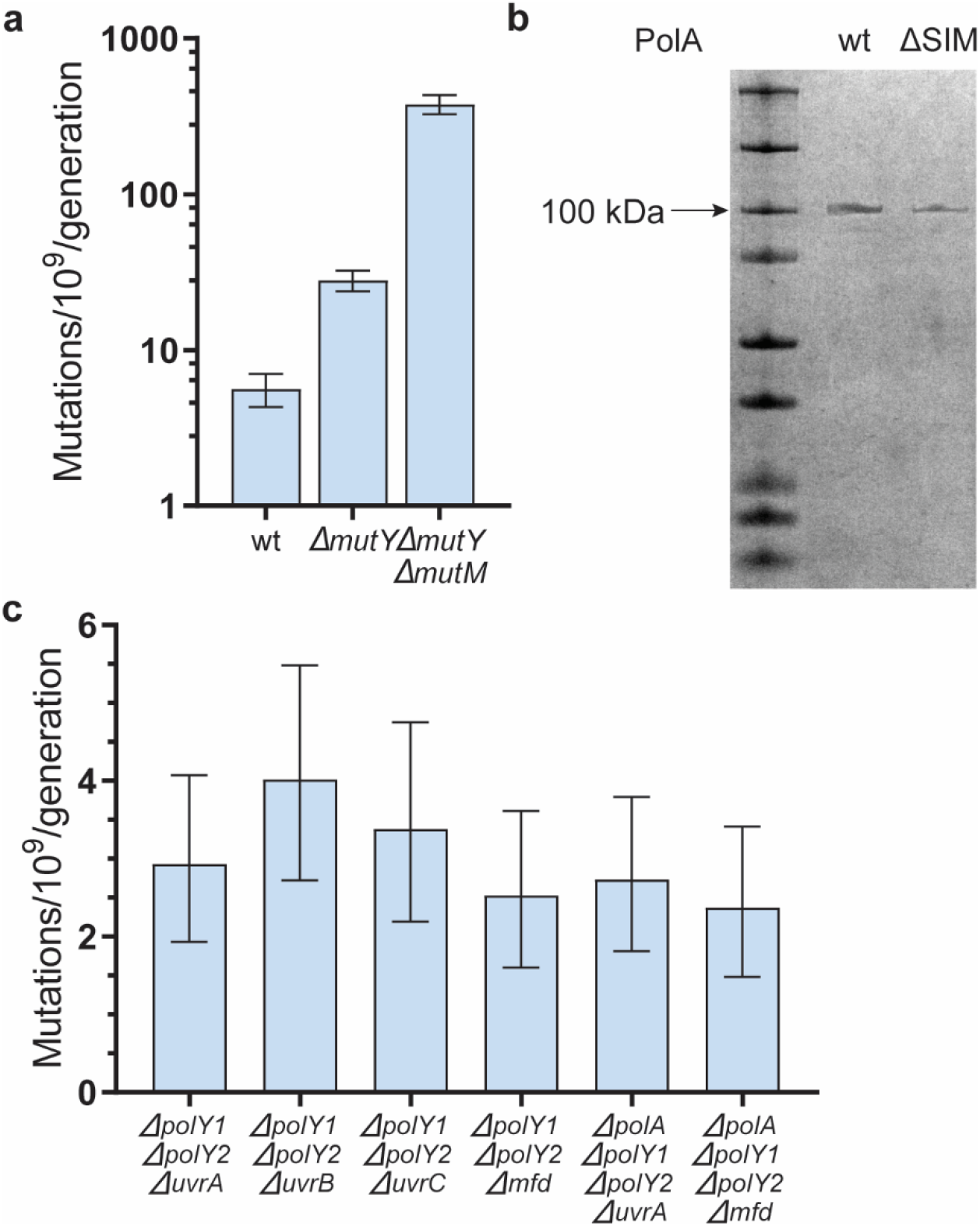
a) Mutation rates of Bacillus subtilis strains of the indicated genotype to rifampicin, n=51 (wt), 59 (*ΔmutY*), 21 (*ΔmutY, ΔmutM*) b) SDS-PAGE of purified *B. subtilis* PolA and PolA-ΔSID. c) Mutation rates of Bacillus subtilis strains of the indicated genotype to rifampicin. n=40 (*ApolY1 ApolY2 AuvrA*), 40 (*ApolY1 ApolY2 AuvrB*), 30 (*ApolY1 ApolY2 AuvrC*), 33 (*ApolY1 ApolY2 Amfd*), 36 (*ApolA ApolY1 ApolY2 AuvrA*), 36 (*ApolA ApolY1 ApolY2 Amfd*) biological replicates. Error bars are 95% confidence intervals.

